# Evaluation of Techniques for Performing Cellular Isolation and Preservation during Microgravity Conditions

**DOI:** 10.1101/040014

**Authors:** Lindsay F. Rizzardi, Hawley Kunz, Kathleen Rubins, Alexander Chouker, Heather Quiriarte, Clarence Sams, Brian E. Crucian, Andrew P. Feinberg

**Affiliations:** Center for Epigenetics, Johns Hopkins University School of Medicine, Baltimore, MD, USA; Science, Technology and Engineering Group, Wyle Laboratories, Houston, TX, USA; Astronaut Office, NASA Johnson Space Center, Houston, TX, USA; Department of Anesthesiology, Hospital of the Ludwig-Maximilians-University, Munich, Germany; JES Tech, Houston, TX, USA; Space and Clinical Operations Division, NASA Johnson Space Center, Houston, TX, USA; Biomedical Research and Environmental Sciences Division, NASA Johnson Space Center, Houston, TX, USA; Departments of Medicine, Biomedical Engineering, and Mental Health, Johns Hopkins University Schools of Medicine, Engineering, and Public Health, Baltimore, MD, USA.

## Abstract

Genomic and epigenomic studies require the precise transfer of small volumes from one container to another. Epigenomic and transcriptional analysis require separation of purified cell types, and long term preservation of cells requires their isolation and transfer into appropriate freezing media. There are currently no protocols for these procedures on the ISS. Currently samples are either frozen as mixed cell populations, with poor yield, or returned under ambient conditions, requiring timing with Soyuz missions. Here, we evaluate the feasibility of translating terrestrial cell purification techniques to the ISS. Our evaluations were performed in microgravity conditions during parabolic atmospheric flight. The pipetting of open liquids in microgravity was evaluated using analog blood fluids and several types of pipette hardware. The best performing pipettes were used to evaluate the pipetting steps required for peripheral blood mononuclear cell (PBMC) isolation via density gradient centrifugation (DGC). Evaluation of actual blood products was performed for both the overlay of diluted blood, and the extraction of isolated PBMCs. We also validated magnetic purification of cells. We found that positive displacement pipettes avoided air bubbles, and the tips allowed the strong surface tension of water, glycerol and blood to maintain a patent meniscus and withstand robust pipetting in microgravity. These procedures will greatly increase the breadth of research that can be performed onboard the ISS, and allow improvised experimentation on extraterrestrial missions.

## Introduction

The ability to collect biosamples onboard the international Space Station (ISS) is extremely limited. With rare exception, only frozen urine, saliva or blood plasma are routinely collected and stored. Recently, the return of ambient blood samples has occurred allowing expanded analyses, but this is constrained to only two samplings per 6-month ISS mission due to availability of the returning Soyuz vehicles. There would be a science benefit for ISS to expand the frequency of flight sampling and to isolate purified cells from blood or tissue while *onboard the ISS,* including peripheral blood mononuclear cells (PBMCs) or positively isolated cell populations (e.g. CD4+, CD8+, or CD19+ cells). This ability would facilitate a host of cellular and ‐omics analyses for which segregation of distinct cell populations has become increasingly important. It is generally perceived that techniques for isolating cells, including density gradient centrifugation (DGC) with Ficoll or magnetic separation of cells, are gravity-dependent. This perception is based on the assumption that open liquids would be difficult to manipulate in microgravity, or that sensitive density-dependent steps such as overlaying of fluids or cellular band removal would be compromised without gravity. We postulated that such steps could be performed in microgravity using standard pipetting techniques. Further, the development of septum-based tubes to facilitate DGC have the potential to render these standard techniques easier in microgravity. These tubes function by the placement of a septum between the Ficoll and blood layers instead of relying on a sensitive overlay, yet with a pore(s) in the septum to allow cellular translocation upon centrifugation. After centrifugation, the septum protects the isolated PBMCs from contamination by red blood cells (RBC) or granulocytes, which would be advantageous on-orbit. We evaluated the fluid handling steps required for DGC, including the loading of Ficoll, the overlay of blood, and the removal and preparation for cryopreservation of isolated cells, in microgravity conditions. Multiple pipetting techniques were evaluated, as were three types (and two sizes) of DGC tube hardware. All evaluations were performed on the NASA C-9 parabolic flight laboratory aircraft. These evaluations demonstrated that the pipetting of open fluids is relatively simple and easily-controlled and that all steps associated with density gradient centrifugation can be replicated in microgravity. Observation of actual purified blood products indicated that the layers created during density gradient centrifugation remain stable in each type of tube evaluated during microgravity. Further, the overlay pipetting of actual diluted blood and the removal of isolated PBMCs was easily accomplished.

## Results

### Testing multiple methods for liquid transfer

We tested five different methods for liquid transfer using several types of pipettes (Figure 1) in microgravity based on several assumptions. The first assumption was that liquids in microgravity would be more susceptible to surface tension effects and consequentially would not necessarily remain in their tubes when opened. To counteract this assumed outcome, we utilized a completely closed system known as a cannula transfer (Supplemental Figure 1) commonly used in chemical synthesis protocols^1^. In order to transfer liquids from one tube to another, the tubes have septa lids that can be punctured by long needles connected to each other via Tygon tubing. Injection of air into the liquid-containing tube increases the pressure in the tube causing the liquid to move up through the needle and tubing into the empty tube. During our initial parabolic evaluations, we discovered that our assumptions were incorrect and that the liquid in the tubes would not spontaneously migrate (Figure 2). The spontaneous movement of fluids, without gravity restraint, generally depends on the wetting characteristics of the fluid, the composition and diameter of the tube itself, and how full the tube is (dependent on the first two conditions). We concluded that the surface tension acting on the liquid in the tubes was sufficient to counteract the reduction in gravity. In addition, the capillary effects were insufficient to cause fluid movement. Nonetheless, we proceeded to test the cannula transfer system. We found that once set up, the transition during the parabola from 1.8 g to microgravity was sufficient to initiate the transfer of liquid through the tubing. While this unassisted transfer would not occur in constant microgravity (as onboard the ISS), the fluid reached a surface tension equilibrium that disrupted our control of the subsequent liquid transfer. We had difficulty forcing the liquid through the tubing after the initial fluid transfer and would occasionally get backflush into the original tube while performing parabolas. Given the complex set up and lack of controlled liquid transfer we eliminated the cannula system as a viable method for transfer.

**Figure 1.**
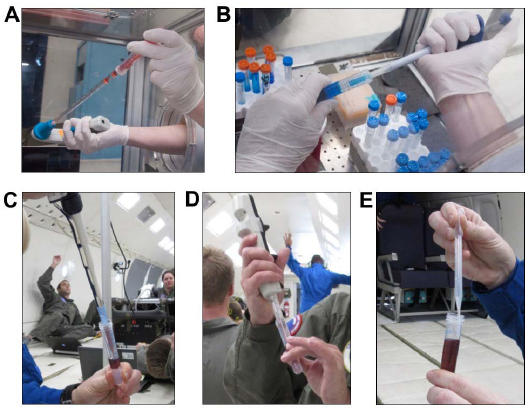
Evaluation of different pipettes for liquid transfer. A) Serological pipette (10 ml) used to transfer dyed water into a 15 ml conical tube. B) Positive displacement pipette (1 ml) used to transfer dyed water into a 15 ml conical tube. C) Standard air displacement laboratory pipette (1 ml) used to transfer dyed water into a 15ml Leucosep tube. D) Positive displacement repeater pipette (5 ml) used to transfer dyed water into a 15ml Leucosep tube. E) Plastic bulb transfer pipette used to transfer dyed water into a 15 ml Leucosep tube.

**Figure 2.**
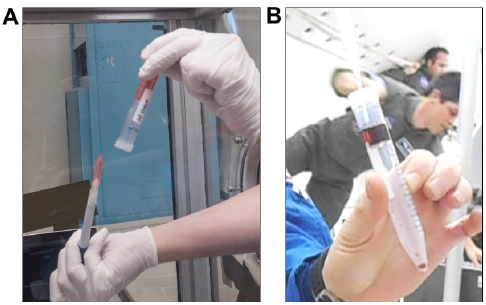
Liquid behavior in microgravity. A) Liquid in the tubes does not spontaneously float out of the tubes, no matter how the tube is oriented. B) Fluids can be easily affixed in place by pipetting anywhere in the tube.

We next tested pipetting with several common laboratory pipettes. Our assumption in this case was that as liquid was drawn up into the pipette tip the reduced gravity would cause 1) the liquid to continue to flow up the inside of the tip and into the pipettor itself (thus contaminating the pipettor) and 2) as a result of this would prevent accurate liquid measurement. The first method we tested utilized plastic serological pipettes (5 and 10 ml capacity) and a battery-operated pipette-aid; there is a cotton plug at the top of the serological pipette (where the pipette enters the pipette-aid) to prevent liquid from being aspirated into the pipette-aid. Once again, our assumptions were unfounded and we had no issues with uncontrolled liquid flowing up the inside of the serological pipette (Figure 1A). However, we found that as we inserted the serological pipette into the tube of liquid we displaced a volume of the liquid that proceeded to migrate up along the inside of the tube. As the pipette was inserted, the wetting, capillary, and surface tension characteristics of the liquid were likely responsible for adherence to the side of the tube and migration along the tube. The migrated liquid would stay at the top of the tube until jostled, at which time the droplet would float away. During the removal process, a small quantity of liquid would sometimes adhere to the pipette itself, and as it was removed a droplet would break off and float away. This significant finding illustrates the importance of the speed of pipette removal as well as accounting for wetting effects. Aside from this issue, we were able to pipette liquid successfully with this method. The difficulty was preventing the introduction of a bubble while doing so, as the migrating liquids altered the liquid level in the tube requiring adjustments to where the pipette tip was placed during the drawing up of the liquid. Bubble formation and release characteristics vary significantly in microgravity compared to terrestrial conditions as has been shown previously^2,3,4^. For these reasons we eliminated this method with the caveat that a larger diameter tube (such that the surface tension is not substantially altered and which allows for less bulk fluid displacement when the serological pipette is inserted) could allow successful use of the serological pipette for transfer.

Next, we tested a plastic bulb transfer pipette (1 ml gradations; 3.1 ml bulb draw). While the least accurate of the pipettes tested, it performed adequately for pipetting water, glycerol, blood and Ficoll analogs, and isolated PBMCs (Figure 1E). A problem arose when liquid was drawn into the bulb portion of the pipette from which it could not be dislodged; however, as long as liquid was drawn up slowly and with control, the transfer pipette was shown to be a viable liquid transfer option. Due to the smaller width of the tip, there were no fluid displacement or surface tension problems as seen with the serological pipets. Together, these results suggest that any tips utilized in microgravity should have a diameter no more than half that of the tubes that will be used (Table 1).

**Table 1.**
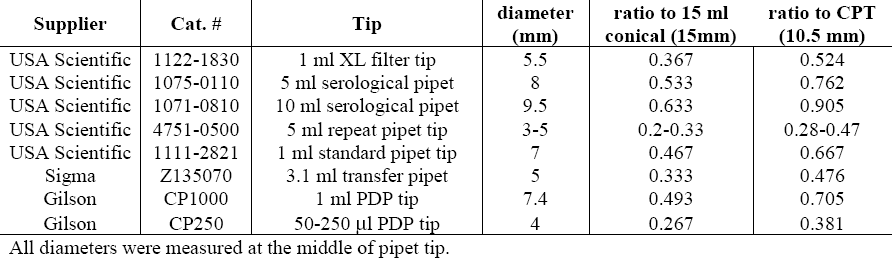
Pipette tip characteristics.

| Supplier | Cat. # | Tip | diameter (mm) | ratio to 15 ml conical (15mm) | ratio to CPT (10.5 mm) |
| --- | --- | --- | --- | --- | --- |
| USA Scientific | 1122-1830 | 1 ml XL filter tip | 5.5 | 0.367 | 0.524 |
| USA Scientific | 1075-0110 | 5 ml serological pipet | 8 | 0.533 | 0.762 |
| USA Scientific | 1071-0810 | 10 ml serological pipet | 9.5 | 0.633 | 0.905 |
| USA Scientific | 4751-0500 | 5 ml repeat pipet tip | 3-5 | 0.2-0.33 | 0.28-0.47 |
| USA Scientific | 1111-2821 | 1 ml standard pipet tip | 7 | 0.467 | 0.667 |
| Sigma | Z135070 | 3.1 ml transfer pipet | 5 | 0.333 | 0.476 |
| Gilson | CP1000 | 1 ml PDP tip | 7.4 | 0.493 | 0.705 |
| Gilson | CP250 | 50-250 µl PDP tip | 4 | 0.267 | 0.381 |

All diameters were measured at the middle of pipet tip.

The final laboratory pipettes tested were a standard air displacement laboratory pipette (1 ml capacity; Figure 1C) and two types of positive displacement pipettes (PDP): an Eppendorf positive displacement repeater pipette (Figure 1D) and Gilson PDPs (capacities ranging from 10 μl – 1 ml) (Figure 1B and Supplemental Video 1) that each use a piston within the tip to allow accurate pipetting while preventing contamination and carry-over. We chose to test this style of pipette in addition to a traditional air-displacement pipette to prevent contamination of the pipette itself in the event of uncontrolled liquid flow. However, our concerns were unfounded as the standard air displacement pipette performed just as well as the PDPs in this regard. These were by far the most successful pipettes tested. We easily and accurately transferred water, analog fluids, glycerol (10% and 100%), and PBMCs using these laboratory pipettes (Figure 1). The only issue we had was again due to the width of the 1 ml Gilson PDP tip relative to the tube diameter when used with the CPT tubes which have a much smaller diameter than the 15 ml conical tubes (Table 1). As long the very end of the pipette tip (smallest diameter) was kept in contact with the liquid this issue was minimized. This issue was not observed with the repeater pipette and lower capacity Gilson PDP tips due to the smaller tip diameters. While it was much less of an issue with the 1 ml PDP than with the serological pipette, in future, we recommend the use of tips with a tip:tube diameter ratio no greater than 0.5 for pipetting in microgravity such as those used with the positive displacement repeater pipette or Gilson PDPs.

### Evaluation of DGC pipetting techniques in microgravity

The steps required for proper execution of a density gradient separation (loading of Ficoll, overlay of sample, and extraction of layer/sample) were evaluated for three types of hardware: Leucosep 15 ml tube, Leucosep 50 ml tube, and a Sepmate 15 ml tube. The 50 ml Leucosep tube is the only one tested that lacks a central pore and is loaded via centrifugation of Ficoll liquid. This fluid loading step could not be performed onboard the aircraft and was performed prior to parabolic flight. For the other pore-containing tubes, it was generally found that the push of fluid through the pore into the confined space for both apparatus was easily performed. Whereas in gravity, a pipetted liquid will ‘fall’ and displace air, in microgravity the introduced liquid intermingled with air, resulting in bubbles (Figure 3A). However, we found that with gentle manual manipulations to push the introduced liquid ‘downward’, we could introduce the necessary volume of analog‐ Ficoll (water and 10% glycerol). A mild centrifugal force was generated by holding the apparatus at the top and gently swinging it outward to create a resultant downward force.

**Figure 3.**
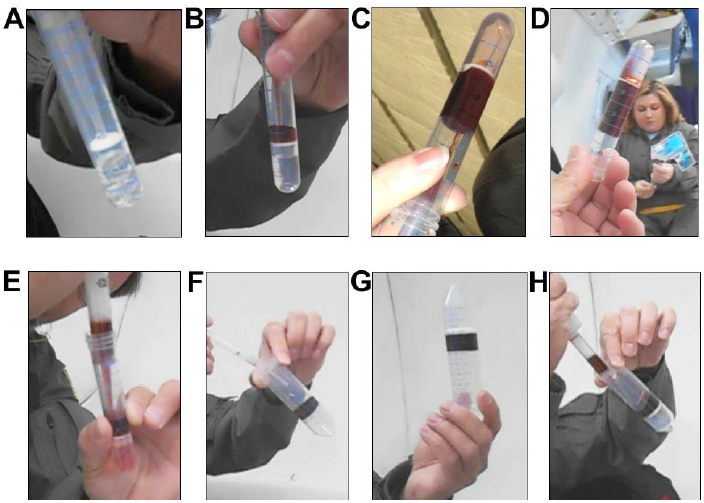
Video capture images documenting evaluation of the steps required to set up a density gradient centrifugation in microgravity conditions. Analog fluids were used: dyed water replaced blood and 10% glycerol in PBS replaced Ficoll. A) Loading of a Leucosep 15 ml tube with Ficoll-analog. B) Overlay of blood‐ analog. C) Demonstration that liquids pipette from up to an inch away are still captured in the appropriate position for the overlay. D) Evidence that mild manipulations did not disturb the integrity of the overlay step. E) Extraction of the overlaid blood analog. F) Overlay of a Leucosep 50 ml tube with blood analog. G) Image demonstrating that when placed in their proper position, the fluids generally remained in place even for a large‐bore 50 ml tube, facilitating operations in microgravity. H) Extraction step for the 50 ml Leucosep tube.

Next, the overlay pipetting step was evaluated for the 15 ml Leucosep and Sepmate DGC apparatus using analog-blood liquid (dyed water) transferred via three pipetting techniques. Note that terrestrially, the overlay is facilitated by gravity, with the introduced fluids being ‘held’ in place over the loaded Ficoll liquid, but even in microgravity, the analog blood was held in place over the analog Ficoll. Fluid was easily aspirated, and then repeatedly pipetted precisely into position creating an acceptable overlay (Figure 3B). We found the overlay was not dependent on carefully ‘placing’ the fluids with the pipette; even pipetting the liquids from approximately 1 inch away allowed capture and collection of the liquid for overlay in the appropriate position (Figure 3C). Further, manipulations of the tube in microgravity did not disturb the overlaid fluids, as evidenced by Figure 3D. Extraction of the overlaid fluids, essentially another aspiration step, which for evaluation purposes was performed without a centrifugation step, was then easily performed (Figure 3E). As the behavior of the open liquids was easily controlled in both 15 ml sized tubes, the protocol was repeated for the larger diameter 50 ml Leucosep tubes. As with the smaller tubes, the overlay of analog-blood was easily performed (Figure 3F). We found it remarkable that even in the larger bore tube, fluids continued to remain in place despite manipulation (Figure 3G). Extraction of the overlaid fluids from the 50 ml tubes was also easily performed in microgravity (Figure 3H). (NOTE: All liquid handling during these evaluations was performed using a standard bulb transfer pipette (3 ml), an Eppendorf positive displacement repeater pipette, and a standard 1000 μl air displacement pipette, each of which performed well.) Video at www.https://youtu.be/5YSeYkCxp4Y provides video support for the results associated with Figure 3.

### Validation of Analog Fluids to Blood Products

The blood samples were processed through the centrifugation step in normal gravity for four types of DGC hardware, but the resulting PBMC ‘band’ (Figure 4A), was not extracted. These processed tubes were then observed during microgravity conditions to evaluate the physical interactions between the blood products and the various tube hardware. The four tubes were a Leucosep 15 ml tube, a Sepmate 15 ml tube, a Becton Dickinson CPT Vacutainer, and a standard 15 ml Falcon tube after normal density gradient separation with Ficoll. The upper layer of the processed samples, consisting of plasma and platelets and containing the PBMC band, remained in place during 3 consecutive parabolic maneuvers which consisted of several altered gravity states (level flight (1 g), 1.8 g, and microgravity) (Figure 4B and 4C) Video at https://youtu.be/xArZFaBLtk provides support for Figure 4. The only deviation was a slight elevation of the meniscus between the plasma and air space within the Leucosep tube indicated by the red arrows in Figure 4B and 4C. After exiting the microgravity period, the meniscus returned to its previous level flight status. These slight alterations did not result in disruption of the PBMC band.

**Figure 4.**
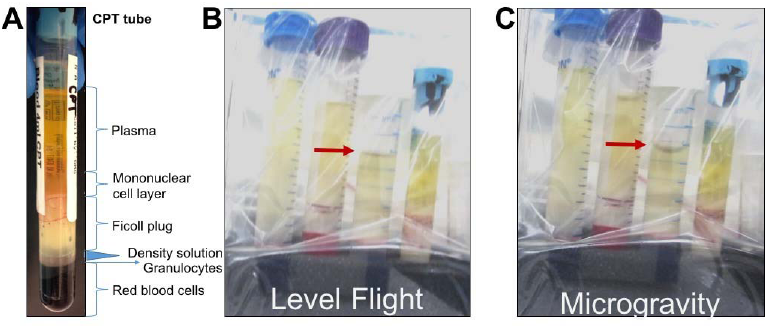
Observation of the behavior of actual blood products processed using density gradient centrifugation during 1 x g level flight versus microgravity conditions. A) Example of processed blood sample in a CPT Vacutainer tube. B and C) Left to Right: Blood products processed and centrifuged via standard Ficoll DGC in a 15 ml Falcon tube, processed and centrifuged using SepMate and Leucosep tubes, and using a Becton Dickinson CPT Vacutainer tube. Tubes were flown triple-contained and video was recorded during several parabolas to ensure blood sample behavior was similar to the analog fluids used in the pipetting evaluation. Note the meniscus of the upper plasma layer (red arrow for the Leucosep 15 ml tubes) is altered during microgravity conditions, but in all tubes the fluids maintained their positions and were never displaced within the tube.

### Blood Evaluation – Overlay Pipetting and Isolating PBMCs in microgravity

To confirm the above findings with actual blood samples, we evaluated overlay pipetting of actual whole EDTA blood diluted 1: 1 with PBS over Ficoll. The Leucosep 15 ml tube was used for this evaluation, with pipetting performed using the 3.1 ml bulb transfer pipette. Blood was aspirated into the transfer pipette, and subsequently overlaid onto the septum of the 15 ml Leucosep tube. Interestingly, we found that there were mild alterations in the behavior of the diluted whole blood. Rather than being easily affixed in place, the blood was slightly ‘sticky’ with a mild affinity for the sidewalls of the tube as evidenced in Figure 5A and 5B. Video at https://youtu.be/xArZFaBLtk provides support for Figure 5. Nevertheless, it was still feasible, with slightly more care than required for the analog-blood fluid, to pipette the blood into place and successfully create the overlay (Figure 5B).

**Figure 5.**
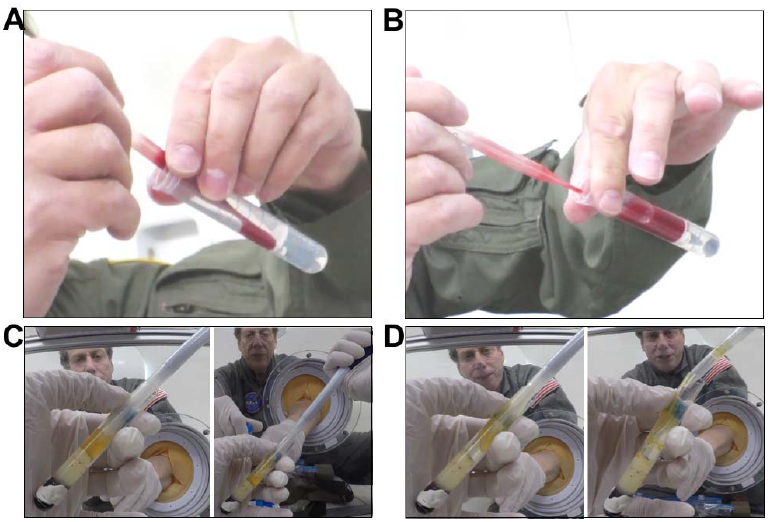
Preparation of a density gradient centrifugation sample using a Leucosep 15 ml tube, loaded with Ficoll reagent overlaid with an actual whole blood sample diluted 1:1 with PBS. The Leucosep tube was preloaded with Ficoll reagent, and a 1:1 diluted whole blood sample was carried aboard in a separate tube. A-B) Using a 3.1 ml bulb transfer pipette, the blood was overlaid onto the device septum in microgravity, effectively yielding a true microgravity preparation of a density gradient separation. C-D) To test the extraction step with actual blood products, PBMCs were extracted during microgravity using a positive displace pipette from CPT Vacutainer tubes prepared prior to parabolic flight. C) Slow and careful pipetting allows for successful aspiration of PBMCs while rapid and careless pipetting (D) makes it impossible to extract precise volumes of PBMCs.

To evaluate the extraction of PBMCs from processed CPT tubes, we collected the blood samples and spun the tubes at Johnson Space Center prior to parabolic flight just as current ISS protocols dictate. The total time from centrifugation to observation in microgravity was approximately 2 hours. As the positive displacement pipettes performed the best, we used the 1 ml capacity PDP to isolate the PBMCs from the CPT Vacutainer tube (Figure 5C and 5D). PBMCs (~ 3 ml) were harvested from the CPT Vacutainer tube by re‐ suspending them in human plasma (top layer in CPT tube; Figure 4A) and transferring them into a 15 ml conical tube containing 333 μl of 100% glycerol for a final glycerol concentration of 10%. Each pipetting step was performed during the microgravity portions of parabolic flight with intervening 1-1.8 g states between. We confirmed in our own lab prior to flight that this concentration of glycerol was sufficient to allow freezing and thawing of viable PBMCs. We observed that re-suspending the cells in the serum as well as mixing with the glycerol can be done without generating bubbles; however, the pipette tip must be kept towards the top of the liquid and mixing via pipetting up and down must be done very slowly to accomplish this (Figure 5C). The CPT tubes are the smallest diameter tubes we tested and if liquid was dispensed too rapidly or the pipette tip displaced too much liquid, bubbles would form or liquid would crawl up the sides of the CPT tube making it very difficult to impossible to precisely aspirate the PBMCs (Figure 5D and Table 1). The PBMC/glycerol mixture was distributed into 3-4 cryovials and stored on ice for the remainder of the flight. After landing, they were transferred to dry ice for transport to Johnson Space Center and stored at ‐80°C.

### Quality control of isolated PBMCs

Microgravity-isolated PBMCs (referred to as ZeroG frozen) were shipped on dry ice to Johns Hopkins for further processing. As a control, we isolated PBMCs in the lab from a different donor via the same protocol used during parabolic flight. One sample was isolated and frozen just as for the microgravity sample (referred to as Hopkins frozen). The other sample was spun in the CPT Vacutainer tube and stored at 4°C overnight to mimic the current protocols used for the NASA Twin Study (referred to as Hopkins ambient). Cells were thawed (when necessary) and total cell number and viability were measured both before and after isolating the CD4+ and CD8+ fractions via magnetic separation with Miltenyi MicroBeads (Table 2). DNA was extracted from each cell type, quantified, and bisulfite converted. Sequencing libraries were generated and checked for quality before destroying the samples. The average library size was 441 bp.

**Table 2.** CD4+ and CD 8+ cell isolation results.

| Sample | Cell Type | Total Cells | Viability | Viable Cells | Total DNA (ng) |
| --- | --- | --- | --- | --- | --- |
| Hopkins Ambient | All Cells | $9 \times 10^6$ | 56% | $5.22 \times 10^6$ | N/A |
| | CD4+ | $1.5 \times 10^6$ | 63% | 945,000 | 702 |
| | CD8+ | $1.1 \times 10^6$ | 72% | 800,000 | 678 |
| | CD4-/CD8- | $5.2 \times 10^6$ | 60% | $3.2 \times 10^6$ | 1890 |
| Hopkins Frozen | All Cells | $5.7 \times 10^6$ | 50% | $2.85 \times 10^6$ | N/A |
|  | CD4+ | 850,000 | 63% | 535,000 | 570 |
|  | CD8+ | 420,000 | 73% | 310,000 | 660 |
| | CD4-/CD8- | $2.4 \times 10^6$ | 33% | 780,000 | 330 |
| ZeroG Frozen | All Cells | $5.7 \times 10^6$ | 45% | $2.56 \times 10^6$ | N/A |
|  | CD4+ | 840,000 | 59% | 495,600 | 882 |
|  | CD8+ | 990,000 | 45% | 450,000 | 672 |
| | CD4-/CD8- | $1.76 \times 10^6$ | 38% | 660,000 | 1038 |

### Evaluation of Magnetic Separation in Microgravity

The basic pipetting steps associated with magnetic cell separation were evaluated during microgravity. Dynabeads suspended in PBS in a 12x75 polystyrene tube were carried onboard the parabolic aircraft (Figure 6A). During microgravity, the magnetic separation was performed by placing the tube in a Dynabead separation magnet (Figure 6B). Separation occurred within 30 sec of microgravity, confirming the assumption that this step would be gravity-independent, as even terrestrially the magnetic force is dominant over gravity, allowing separation in a typical laboratory setting. It was observed that bubbles would permeate the liquid and not ‘rise’ and thus be eliminated from the system (Figure 6B). However, the presence of bubbles did not seem to inhibit the separation and a proper isolation of the magnetic beads on the tube sidewall. Extraction of the liquid phase, required for subsequent cell purification, was easily performed with either a bulb transfer pipette (Figure 6C) or a standard 1000 μl air displacement laboratory pipette (Figure 6D). Video at https://youtu.be/UZINFN0gERQ depicts the steps associated with magnetic separation performed in microgravity conditions.

**Figure 6.**
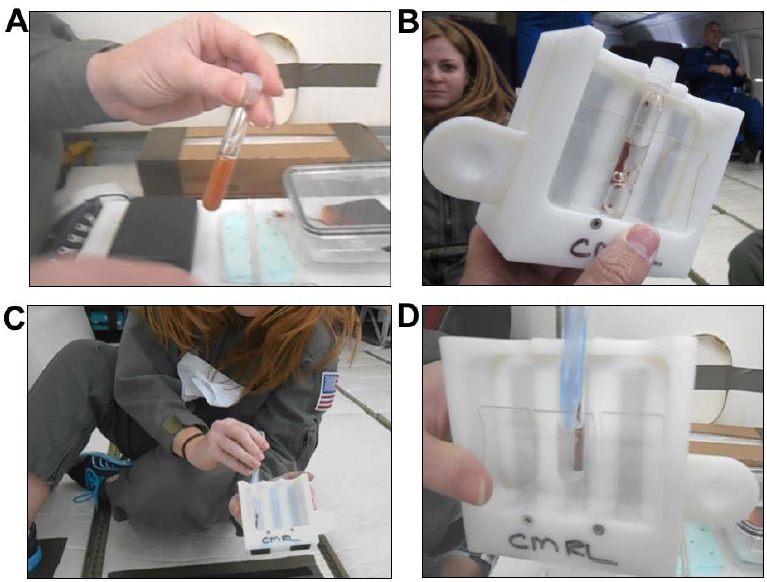
Evaluation of magnetic separation using Dynabeads,. A) Dynabeads suspended in PBS. B) Magnetic separation of beads using a Dynabead separation magnet. C and D) Extraction of the liquid phase without disturbing the magnetic beads using a transfer pipette (C) .and a standard air displacement pipette (D).

## Methods

### Biological and analog samples

The microgravity evaluation was primarily performed using analog fluids. Dyed water represented blood, and 10% glycerol in 1X PBS represented Ficoll solution. 100% glycerol was also tested to determine how very viscous fluids would perform in microgravity. EDTA blood was collected from a single healthy test subject. Prior to flight, the EDTA sample was diluted 1:1 with isotonic 1X PBS. Ficoll (Sigma, St. Louis, MO) was flown unaltered. Blood from two additional healthy test subjects was collected into CPT Vacutainer tubes. The NASA laboratory maintains an IRB approval for blood sample collection related to assay development and instrument validation, including microgravity testing, which applied to this activity.

### Evaluation hardware

Three types of hardware which support density gradient separation were evaluated during microgravity conditions: Leucosep 15 ml, Leucosep 50 ml, and Sepmate 15 ml tubes. Each of these tubes possesses a ‘septa’ which separates the Ficoll from the sample (prior to centrifugation) and isolates the PBMC/plasma from the RBC/granulocytes (after centrifugation). Five types of pipettes were used: a standard bulb transfer pipette (1 ml gradations; 3.1 ml bulb draw), serological pipettes with battery-powered pipette-aid (5 and 10 ml), an Eppendorf positive displacement repeater pipette, a series of Gilson (Middleton, WI) positive displacement pipettes (10 μl – 1000 μl), and a standard 1000 μl (blue tip) pipette. We also tested a cannula transfer system that consisted of 15 ml conical tubes with septa lids that can be punctured by long needles (18G, 3.5 inch Tuohy Epidural needles) connected to each other via luer-locks and Tygon tubing. One 15 ml tube contained dyed water while the other was empty. Liquid transfer was initiated by injecting air into the water-filled tube via a standard 10 cc syringe and 25G needle. We also tested performance of magnetic Dynabeads (Thermofisher) along with the Dynabead magnet (Thermofisher) with blood/Ficoll analogs.

### Microgravity conditions

NASA can perform actual microgravity evaluation of hardware on Earth by using parabolic flight aircraft. A NASA C-9 jet aircraft flies in a parabolic arc, where an approximately 30 sec period of microgravity is generated at the top of the parabola, and a corresponding period of 1.8 g is generated at the parabola bottom. On a flight, 40 parabolas are flown in a repeated sequence, resulting in forty ~ 30 sec microgravity opportunities. For this evaluation, four flight days were utilized, and all steps were performed within the ~ 30 sec of microgravity. Flight logs available upon request.

### Protocol for transferring PBMCs from CPT Vacutainer tube in microgravity

Blood was drawn into a CPT Vacutainer tube, inverted 10 times, incubated at room temperature (RT) for 15 min, and spun at 1800 x g for 20 min at RT to separate the PBMC layer (Figure 4A). The CPT tube was then taken onto the aircraft for the microgravity parabolas (approximate 2 h delay). All the sample handling was done in microgravity (with the on-board gravity indicator at zero) with pauses at each gravity transition (from 1.8 g to microgravity). While working inside a glove box, a 1 ml PDP was used to transfer the topmost 1 ml of serum into a 15 ml conical tube. After the next g transition, we mixed the remaining serum carefully by pipetting to re-suspend the cell layer while in microgravity. We removed 1 ml at a time until less than 1 ml remained. Due to the low volume and width of the pipette tip we had to use a transfer pipette for the remainder; this would be unnecessary if we had used the repeater PDP which has a much narrower tip (Table 1). We estimated the volume from the incremental measurements displayed on the tube. We added 100% glycerol to a final concentration of 10% using an appropriately sized PDP. We pipetted up and down carefully to thoroughly mix, transferred 900 μl into each of three cryovials and placed on ice (all pipetting occurred during the microgravity portions of parabolic flight). Once they had been mixed, we did not observe any separation of the glycerol and cell mixture during g transitions. Upon landing, the tubes were transferred to dry ice and then to an insulated container to freeze at ‐80°C.

### Cell type isolation at Johns Hopkins using Miltenyi magnetic beads

Frozen aliquots of PBMCs (Hopkins frozen and microgravity samples) were thawed rapidly in a 37°C water bath, combined in a 15 ml conical tube (if from the same sample), and washed twice with 10 ml of 1X phosphate buffered saline (PBS), pH 7.5. PBMCs isolated from the Hopkins ambient sample were also washed twice in 10 ml 1X PBS. Cells were collected by centrifugation at 300 x g for 10 min at 4°C and resuspended in 5 ml of cold Cell Suspension Buffer (CSB; 1X PBS containing 2 mM EDTA, 0.5% BSA). Total cell count and viability were measured using the Countess Automated Cell Counter (Invitrogen, Carlsbad, CA, USA). Cells were spun again at 300 x g for 10 min at 4°C and resuspended in 80 μl CSB. To isolate CD4+ cells, 20 μl of CD4+ magnetic beads (Miltenyi, Auburn, CA) was added, mixed by pipetting, and incubated 15 min on ice in the dark. Following the incubation, 5 ml of cold CSB was added and the samples were spun at 300 x g for 10 min at 4°C. To capture the CD4+ cells, the pellet was resuspended in 1 ml CSB and passed through the MS columns attached to a magnet stand (Miltenyi, Auburn, CA) as per the manufacturer’s instructions. The flow through was saved and used for the isolation of CD8+ cells following the same protocol, but using the CD8+ magnetic beads. Once the CD4+ cells were captured on the column, a 500 μl CSB wash was performed. To elute the CD4+ cells into a 1.5 ml Eppendorf tube, the column was removed from the magnet, 1 ml of CSB was added, and the included plunger was used to expel the CD4+ cells. Total cell and viability counts for the CD4+, CD8+ and the remaining unlabeled cell fraction were performed using the Countess Automated Cell Counter (Invitrogen, Carlsbad, CA, USA). The cells were pelleted by centrifugation at 300 x g for 10 min at 4°C and DNA was isolated using the MasterPure kit (Epicentre, San Diego, CA). Total DNA was quantified using the Qubit dsDNA BR Assay kit (Invitrogen, Carlsbad, CA, USA). The DNA was stored at ‐20°C until use.

### Bisulfite conversion and library prep

Whole genome bisulfite sequencing (WGBS) single indexed libraries were generated using NEBNext Ultra DNA library Prep kit for Illumina (New England BioLabs, Ipswich, MA, USA) according to the manufacturer's instructions with modifications. Five hundred nanograms of gDNA was quantified by Qubit dsDNA BR assay (Invitrogen, Carlsbad, CA, USA) and fragmented by Covaris S2 sonicator to an average insert size of 350 bp. Size selection was performed using AMPure XP beads and insert sizes of 300-400 bp were isolated. Samples were bisulfite converted after size selection using the EZ DNA Methylation-Gold Kit (Zymo, Irvine, CA, USA) following the manufacturer’s instructions. Amplification was performed after the bisulfite conversion using Kapa Hifi Uracil+ (Kapa Biosystems, Boston, USA) polymerase using following cycling conditions: 98°C 45 sec /8cycles: 98°C 15 sec, 65°C 30 sec, 72°C 30 sec / 72°C 1 min.

Final libraries were run on 2100 Bioanalyzer (Agilent, Santa Clare, CA, USA) High-Sensitivity DNA assay, samples were also run on Bioanalyzer after shearing and size selection for quality control purposes. Libraries were quantified by qPCR using the Library Quantification Kit for Illumina sequencing platforms (KAPA Biosystems, Boston, USA), using the 7900HT Real Time PCR System (Applied Biosystems).

## Discussion

Although common perception is that handling of open liquids would be difficult in microgravity, we found that all pipetting steps related to density gradient centrifugation were easily performed in microgravity using analog fluids as well as human blood products (Figure 1). We previously observed that glass and plastic blood collection tubes fill differently in microgravity. In plastic tubes blood tends to fill ‘top to bottom’ in any orientation, whereas blood has a greater affinity for glass vacutainers. In glass tubes, blood tends to rapidly ‘coat’ the interior surface, resulting in a hollow ‘core’ of air until the tube completely fills (authors’ unpublished observations from parabolic flight and spaceflight onboard ISS). This observation in part contributed to our assumption that liquid handling (particularly of actual blood) would be quite difficult during microgravity conditions. However, we found that processed blood products and Ficoll behaved similarly to their analogs (water and glycerol) and could easily and accurately be manipulated via pipetting (Figure 5 compared to Figure 2).

We tested a variety of pipetting mechanisms and found that the most important factor for success was the ratio of diameters between the pipet/tip and the liquid-containing tube. The air-displacement, positive‐displacement, and transfer pipettes performed the best for each liquid and tube type tested as they had the smallest tip diameters compared to the larger serological pipets (Table 1). At no point during microgravity did liquid spontaneously flow out of the tubes as we initially expected (Figure 2). The most difficult aspect of liquid handling proved to be preventing the introduction of bubbles into the liquids during transfer or mixing. Once bubbles were introduced, there was no way to remove them or to pipet around them during microgravity. This finding underscores the absolute requirement for very careful and slow pipetting to ensure bubbles are never introduced.

After determining that the PDPs and standard laboratory air displacement pipettes are the most accurate and easy-to-use methods of liquid transfer, we have a few recommendations to further simplify liquid transfer using this method. The first is to utilize a 5 ml capacity pipettor (e.g. ErgoOne 5 ml pipettor; cat#: 7150-5000) which would prevent multiple pipetting steps for larger volumes (such as in the CPT tubes), decreasing the opportunities for introducing bubbles. Care would need to be taken to insert only the end of the tip (2 mm diameter) as the diameter increases to 11 mm towards the middle of the pipette tip. Ideally, smaller diameter tips could be manufactured for this pipettor to prevent the fluid displacement and surface tension issues we demonstrated in microgravity. For smaller volumes, the currently available PDPs and air-displacement pipettors (ranging in capacity from 10 μl to 1 ml) perform well. Secondly, a “space-ready” gripper box for the pipette tips would allow the box to remain open while holding the tips in place (Supplemental Video 2). A tube holder that allows the user to snap various sizes of tubes in place (perhaps using a snap clamp such as those used for vortex mixers) would also be helpful.

This study has found that all steps needed for DGC and cell isolation can be completed in microgravity conditions. The deployment of a swinging bucket centrifuge to ISS (currently anticipated in 2017) will be the final necessary hardware to allow purified mononuclear cells to be easily isolated onboard ISS. The steps required for cell isolation via DGC (Figure 5) and magnetic bead separation of cells (Figure 6; both pipetting and magnetic ‘pull’) were demonstrated during this parabolic flight evaluation. Coupling DGC with magnetic separation would allow the isolation of purified types of peripheral leukocytes onboard ISS. This capability would greatly enhance onboard sample collection for various types of ‐omics analyses, with the potential to complement existing biosample repository collections.

## Acknowledgements

The authors would like to thank Jerry Myers and John Mcquillen (NASA Glenn Research Center) for advice and critical reading of the manuscript. We also thank Terry Lee, Kerry McMannis, D. Del Rosso and the pilots and other crew members from NASA’s Reduced Gravity Office for coordinating our use of the C-9 aircraft for parabolic flights. We thank NASA photographers/videographers James Blair and Regan Geeseman for documenting our work. Additionally, we thank Veronica Seyl for use of the Education Office’s glove box for these experiments as well as Marisa Covington and everyone at Johnson Space Center for their contributions to the success of this work. This work was supported by funding from NASA (NNX14AH28G awarded to APF and a Division Innovation award from the NASA Biomedical Research and Environmental Sciences Division to BEC).

### Contributions

LFR, KR, CS, BC, and APF, designed the study. Experiments were performed by LFR, HK, KR, BC, and APF. LFR, HK and BC wrote the paper with input and editing from all authors.

### Competing Interests

The authors declare no conflict of interest.

---

**Supplemental Figure 1.**
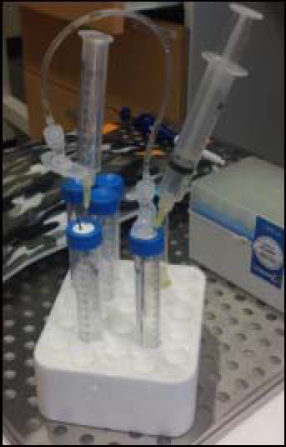
Terrestrial set up of cannula transfer. Tygon tubing attached to two needles connects two 15 ml conical tubes with and without liquid. A syringe is used to inject air into the liquid containing tube to force the liquid through the tubing into the empty tube. A syringe without a plunger is used to vent the displaced air from the empty tube as liquid enters.

**Supplemental Video 1.** Video of pipetting colored water using a Gilson 1 ml PDP in microgravity. Demonstrates the ease of liquid transfer and stability of the liquid in the tube.

**Supplemental Video 2.** Video demonstrating need for a “space-ready” gripper box for pipette tips as they will float up out of the standard tip box when lid is left open.

